# MARCH family E3 ubiquitin ligases selectively target and degrade cadherin family proteins

**DOI:** 10.1101/2023.08.10.552739

**Authors:** Tadahiko Seo, Anthony M. Lowery, Haifang Xu, William Giang, Sergey M. Troyanovsky, Peter A. Vincent, Andrew P. Kowalczyk

## Abstract

Cadherin family proteins play a central role in epithelial and endothelial cell-cell adhesion. The dynamic regulation of cell adhesion is achieved in part through endocytic membrane trafficking pathways that modulate cadherin cell surface levels. Here, we define the role for various MARCH family ubiquitin ligases in the regulation of cadherin degradation. We find that MARCH2 selectively downregulates VE-cadherin, resulting in loss of adherens junction proteins at cell borders and a loss of endothelial barrier function. Interestingly, N-cadherin is refractory to MARCH ligase expression, demonstrating that different classical cadherin family proteins are differentially regulated by MARCH family ligases. Using chimeric cadherins, we find that the specificity of different MARCH family ligases for different cadherins is conferred by the cadherin transmembrane domain. Further, juxta-membrane lysine residues are required for cadherin degradation by MARCH proteins. These findings expand our understanding of cadherin regulation and highlight a new role for mammalian MARCH family ubiquitin ligases in differentially regulating cadherin turnover.

## Introduction

Cadherin family cell adhesion molecules play key roles in a wide range of developmental processes and in pathologies such as tumorigenesis [1-3]. Cadherins are particularly important in the establishment of epithelial and endothelial barriers in the skin, vascular system, and other organs [4-6]. In a number of disease contexts, cadherin expression levels are reduced, or the type of cadherin expressed is altered. For example, in Kaposi sarcoma, an endothelial derived tumor, VE-cadherin is downregulated [7-9]. Likewise, E-cadherin is often down-regulated in a variety of epithelial tumors, including pancreatic carcinoma, gastric cancer, and breast cancer [10-12] . On the other hand, N-cadherin is upregulated in various tumors and its expression is thought to enhance cell migration [13-15]. Indeed, upregulation of N-cadherin at the expense of other classical cadherins is a hallmark of endothelial and epithelial to mesenchymal transitions in developmental processes and tumor progression.

A key regulatory mechanism for cadherin cell surface expression is down-regulation by endocytic membrane trafficking pathways [16-18]. While β-catenin functions primarily to couple cadherins to the actin cytoskeleton, p120-catenin functions as a set point for cadherin levels by inhibiting cadherin endocytosis [19, 20]. In the case of VE-cadherin, p120-catenin binds to and masks a constitutive endocytic motif in the cadherin cytoplasmic tail comprising three acidic residues DEE [21]. This DEE motif mediates clathrin-dependent endocytosis of VE-cadherin and is highly conserved in other classical cadherins across species [21]. p120-catenin gene ablation in mice [22] causes lethality, and mutation of the p120-catenin binding site in VE-cadherin [23] causes vascular leak. Mutation of the DEE endocytic motif compromises the establishment of endothelial cell polarity during migration and leads to angiogenic defects [23]. Thus, VE-cadherin endocytic regulation is essential for the development of a functional vascular barrier and for normal angiogenic patterning of vessels.

In addition to the VE-cadherin DEE sequence, we and others have also shown that cadherin cell surface levels can be down regulated by ubiquitin dependent pathways [7, 9, 24]. K5 is a ubiquitin ligase present in the human herpes virus 8 (HHV-8) genome that downregulates VE-cadherin through ubiquitination of two membrane proximal lysine residues [9]. This process is thought to be important in the development of Kaposi sarcoma vascular tumors that are frequently observed in immune-suppressed individuals infected with HHV-8. Interestingly, K5 exhibits homology to human Membrane-Associated RING-CH-type ubiquitin ligases (MARCH) [25]. The MARCH family proteins are known to affect immune system related proteins such as MHC1 and MHC2 [25-27]. Interestingly, MARCH8 downregulates E-cadherin, and knockdown of MARCH8 causes developmental defects in zebrafish embryos [28]. Similarly, MARCH3 has been reported to downregulate VE-cadherin, but indirectly, through a transcriptional regulator [29]. However, the contribution of MARCH family proteins to cell adhesion is poorly understood, and the mechanism by which MARCH family ligases select cadherins for degradation is unknown.

In the present study, we sought to determine which MARCH family ligases target individual cadherins and how selectivity for cadherin degradation is achieved. Our results indicate that MARCH family proteins differentially downregulate cadherin family proteins with specificity that is conferred by the cadherin transmembrane domain. These findings have important implications for understanding how the cadherin repertoire of various cell types might be regulated in the context of vertebrate development and tumorigenesis.

## Results and discussion

### MARCH family ligases exhibit selectivity for different cadherins

Based on the known role of the Kaposi sarcoma ligase K5 in downregulating endothelial cadherins, we initiated studies using endothelial cells to determine how MARCH family ligases impact various cadherin levels. The expression profile of MARCH family genes in human endothelial cells was assessed by RT-PCR of HUVEC lysates. Human endothelial cells express MARCH2, 3, 4, 5, 6 and 8 (Fig 1A). We focused on MARCH2, 3, 4 and 8 due to their structural relationship with the HHV-8 ligase K5, their expression profile, and based on previous studies linking some of these ligases to cadherin turnover [9, 28]. GFP-tagged MARCH2, 3, 4 and 8 were transfected into HUVECs, and the localization of endogenous endothelial cadherins VE-cadherin and N-cadherin was assessed by immunofluorescence (Figs 1B and C). Expression of MARCH2-GFP and MARCH3-GFP decreased VE-cadherin localization at cell borders but had no discernible effect on N-cadherin (Figs 1B and C). Expression of MARCH8-GFP slightly downregulated VE-cadherin but not N-cadherin (Figs 1B and C), whereas MARCH4-GFP had no effect on either VE-cadherin or N-cadherin (Figs 1B and C). We also tested the effect of MARCH family proteins on E-cadherin, which plays a central role in epithelial adherens junctions. E-cadherin was downregulated by MARCH8, as previously reported [28], but was not altered by other MARCH family proteins (Fig S1). p120-catenin is a well-known regulator of classical cadherin cell surface levels [30, 31]. Therefore, we also examined the localization of p120-catenin by immunofluorescence. p120-catenin remained at cell borders in the presence of all MARCH ligases tested with the exception of MARCH2. Cells expressing MARCH2-GFP exhibited some reduction in p120-catenin levels at cell-cell contacts (Fig 1D). These results suggest that different MARCH family ligases directly target different cadherin family members, with MARCH2 and MARCH3 selective for VE-cadherin. Surprisingly, N-cadherin was uniquely resistant to all MARCH ligases tested.

**Fig 1.**
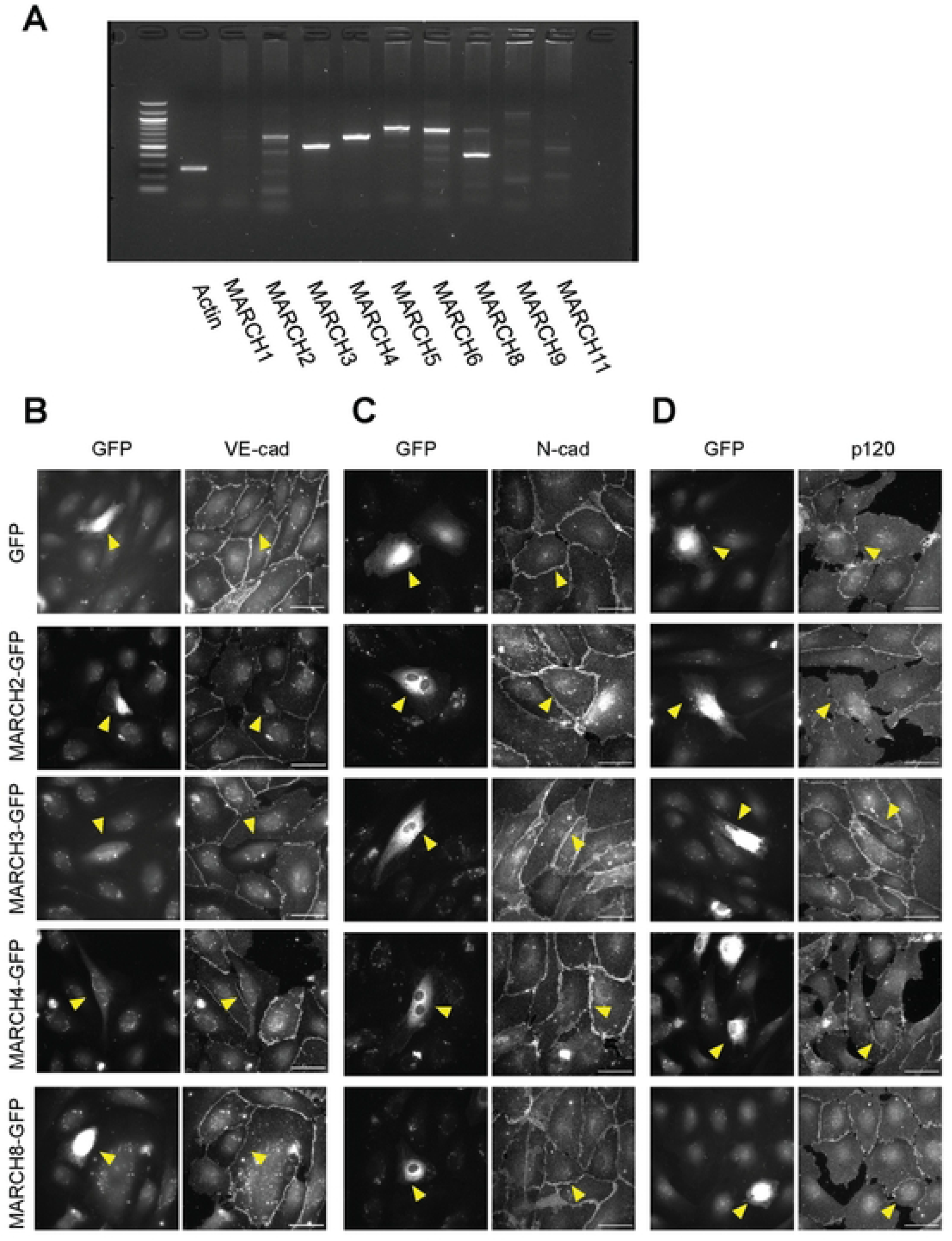
MARCH family ubiquitin ligases exhibit differential abilities to down-regulate various cadherin family proteins. (A) RT-PCR for MARCH family ligases in HUVECs. (B, C, D) Endothelial cells were transfected for GFP alone or with various GFP-tagged MARCH family proteins. Cells were incubated 24 hours and then immunostained for VE-cadherin (B), N-cadherin (C) or p120-catenin (D). GFP positive cells are indicated by yellow arrowhead. Scale bar = 50 µm

### MARCH2 compromises endothelial barrier function

To verify that MARCH ligase activity was required for loss of junctional VE-cadherin, a ligase dead MARCH2 variant (MARCH2-LD-GFP) was constructed. Consistent with the results in Fig 1, cells expressing MARCH2-GFP lost VE-cadherin but not N-cadherin (Fig. 2A-C). In contrast, the MARCH2-LD-GFP mutant and MARCH4-GFP did not alter VE-cadherin localization (Fig 2A). p120-catenin accumulation at cell borders decreased in cells expressing MARCH2-GFP (Fig 2B) but overall p120-catenin levels remained similar in each group as assessed by western blot analysis (data not shown). Interestingly, N-cadherin expression increased in MARCH2-GFP expressing cells but not in MARCH2-LD-GFP or MARCH4-GFP expressing cells (Figs 2B and C). The increase in N-cadherin likely reflects an increase in availability of p120-catenin binding to N-cadherin when VE-cadherin levels are down-regulated, as previously reported [32]. Interestingly, the organization of the actin cytoskeleton was dramatically altered by MARCH2-GFP transfection but not by MARCH2-LD-GFP or MARCH4-GFP (Fig 2A). Because VE-cadherin is thought to modulate VEGFR signaling, we also assessed VEGFR2 levels [33-35]. Interestingly, MARCH2 but not MARCH4 increased VEGFR2 levels. Lastly, we assessed whether MARCH2 affects the integrity of the endothelial cell barrier using electric cell-substrate impedance sensing (ECIS). MARCH2-GFP transduction significantly impaired vascular barrier integrity in a manner dependent upon ligase activity (Fig 2D). Together, these results indicate that MARCH2 specifically downregulates VE-cadherin and compromises endothelial barrier integrity in a ubiquitin ligase dependent manner. Furthermore, the presence of N-cadherin is insufficient to maintain endothelial barrier function.

**Fig 2.**
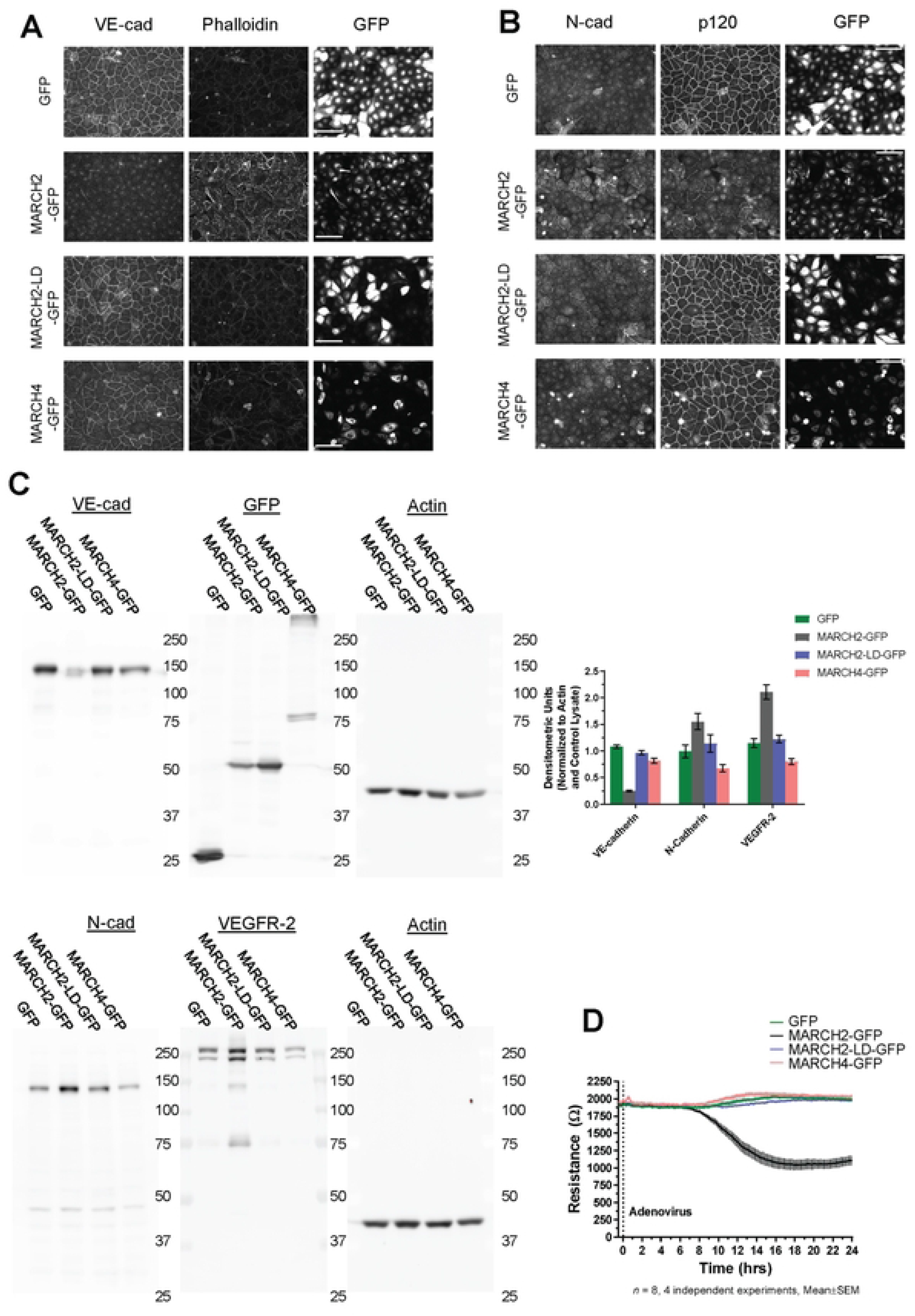
MARCH2 compromises endothelial cell-cell junctions and reduces endothelial barrier function. Confluent monolayers of primary human umbilical vein endothelial cells (HUVEC) were infected with adenoviral vectors to express MARCH2-GFP, MARCH2 LD-GFP, MARCH4-GFP or GFP. After 24 hours cells were fixed for immunofluorescence or lysed for western blot analysis. Monolayers were co-labeled for (A) VE-cadherin and F-actin or (B) N-cadherin and p120-catenin (Scale bar = 100 µm). Expression of GFP or GFP tagged MARCH ligases were also visualized by fluorescence in these monolayers. (C) Western blot analysis for VE-cad, N-cad, VEGR-2, GFP and actin. Band density for VE-cad, N-cad, VEGFR-2, and actin were quantified by Multi Gauge (V3.0) and presented as mean± sem, n= 4 (D) Confluent monolayers grown on ECIS chamber slides were infected with adenoviral vectors, and trans-endothelial electrical resistance was continuously measured by ECIS for 24 hrs.

### The transmembrane domain of VE-cad is necessary and sufficient for MARCH2 specificity

We next sought to determine the basis for cadherin selectivity by MARCH family ligases. Classical cadherins contain endocytic motifs nestled within the p120-catenin binding site on the cadherin juxtamembrane domain [21, 36, 37]. Previous studies have implicated transmembrane domains in MARCH ligase substrate selectivity and found that the ligases typically target membrane proximal lysine residues [38-40]. Therefore, we constructed chimeric cadherins by swapping transmembrane domains and juxtamembrane domains of VE-cadherin and N-cadherin. We generated four chimeras: VE-cadherin with a N-cadherin transmembrane domain (VE-cad-NcadTMD-RFP), VE-cadherin with N-cadherin transmembrane and juxtamembrane domains (VE-cad-NcadTMDJMD-RFP), N-cadherin with a VE-cadherin transmembrane domain (N-cad-VEcadTMD-RFP), and N-cadherin with VE-cadherin transmembrane and juxtamembrane domains (N-cad-VEcadTMDJMD-RFP) (Fig 3A). The various constructs were assembled into a lentivirus delivery system and used to generate stable A431 epithelial cell lines. A431 cells were chosen as these cells lack both VE-cadherin and N-cadherin (Fig S3 and [41]). As shown in Fig S2, all of the chimeric cadherins were uniformly expressed and assembled at cell-cell contacts similarly to wild type cadherins.

**Fig 3.**
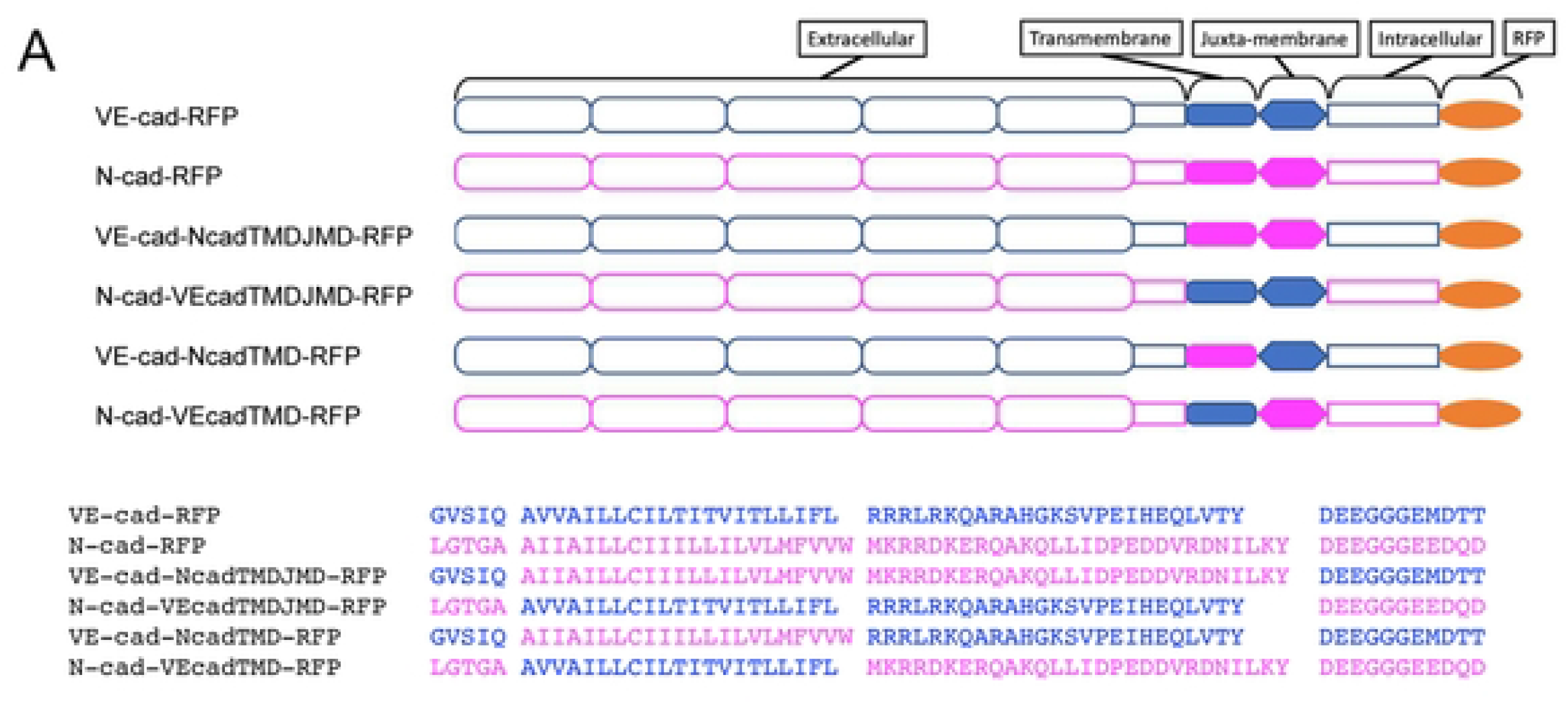
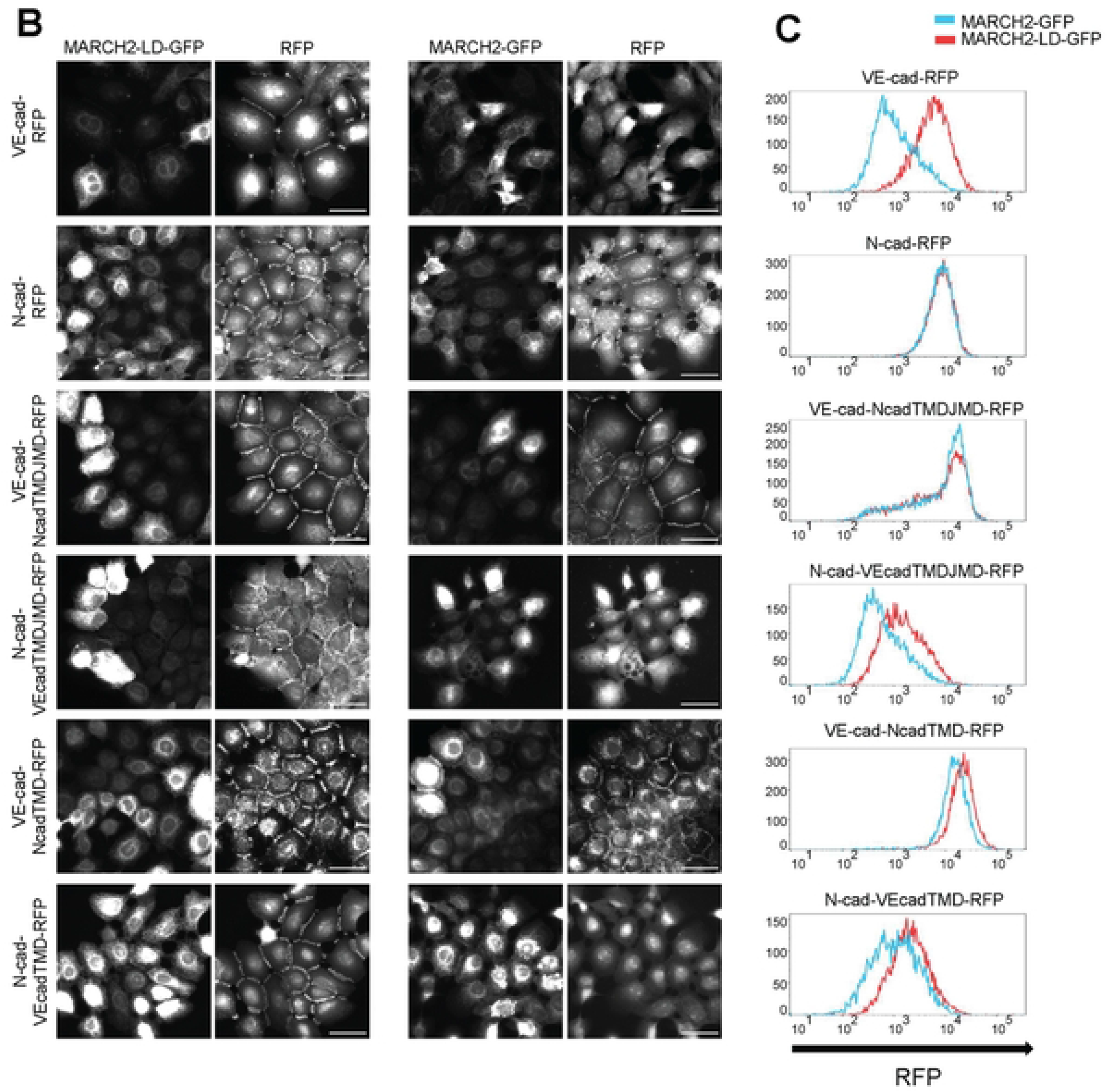
The cadherin juxta-membrane and transmembrane domains of VE-cad specify sensitivity to MARCH2-dependent degradation. (A) Cadherin domains and chimeras are indicated by schematic. VE-cadherin derived parts are color-coded blue while N-cadherin derived parts are magenta. The amino-acid sequence of transmembrane and juxta-membrane domains of the cadherin constructs are indicated below the schematics (B) MARCH2-LD-GFP (left) and MARCH2-GFP (right) were expressed in A431 cells expressing the various cadherin chimeras using an adenoviral expression system. 24 hours after transduction, cells were fixed and processed for fluorescence microscopy. Scale bar = 50 µm (C) Flow cytometry for the cadherin chimeras in cells expressing MARCH2-LD-GFP or MARCH2-GFP. Living GFP positive cells were acquired by filtering forward scatter (FSC), side scatter (SSC), and GFP values. Vertical axis indicates counts of cells, and horizontal axis indicates cadherin-RFP signal intensity. Red line indicates MARCH2-LD-GFP transfected cells, and blue line indicates MARCH2-GFP transfected cells.

To determine which cadherin domain was required for MARCH2 sensitivity, cells expressing the chimeric cadherins were transduced with adenovirus to express either MARCH2-GFP or MARCH2-LD-GFP. Fluorescence localization analysis confirmed expression of the ligases and cadherins (Fig 3B). To quantitatively compare cadherin levels in cells expressing similar levels of the MARCH ligase, we used flow cytometry to monitor MARCH2 expression cadherin levels using GFP and RFP respectively. This approach allowed us to quantitatively compare cadherin levels in cells expressing similar levels of the MARCH ligase. Consistent with the results observed in endothelial cells, MARCH2 downregulated wild type VE-cadherin but not N-cadherin. Interestingly, when the N-cadherin transmembrane and juxtamembrane domains were replaced with the corresponding domains from VE-cadherin (N-cad-VEcadTMDJMD and N-cad-VEcadTMD), the resulting N-cadherin chimera became sensitive to MARCH2 down-regulation. Furthermore, when the opposite approach was taken and the VE-cadherin TMD and JMD were replaced by corresponding sequences from N-cadherin (VE-cad-NcadTMDJMD and VE-cad-NcadTMD), the chimeras were largely resistant to down-regulation by MARCH2 (Figs 3B and C). These results demonstrate that the cadherin transmembrane domains are essential for conferring selective recognition by MARCH family ligases. These findings are consistent with previous studies for other MARCH family substrates linking the transmembrane domain to substrate recognition by MARCH family ligases [40].

### The downregulation of VE-cadherin by MARCH2 depends on juxtamembrane lysine residues but occurs independently of the DEE constitutive endocytosis motif

We previously showed that the HHV-8 ubiquitin ligase K5 targeted two lysine residues in the juxtamembrane of VE-cadherin for ubiquitin mediated endocytosis and degradation [9]. However, VE-cadherin also harbors a unique endocytic motif (DEE) in the juxtamembrane that also functions as a p120-catenin binding site [21]. To investigate which pathway mediates MARCH2-induced VE-cadherin downregulation, we constructed RFP-tagged variants harboring lysine to arginine mutations (VE-KK-RFP), mutations in the DEE constitutive endocytic motif (VE-DEE-RFP), or mutations in both the ubiquitin ligase and constitutive endocytic motifs (VE-KKDEE-RFP) (Fig 4A). These VE-cadherin mutants assembled into adherens junctions similarly to wild type VE-cadherin, consistent with previous results examining the localization of E-cadherin endocytic mutants [42]. We then assessed the effect of MARCH2 on each mutant (Figs 4B-E). MARCH2 transduction caused VE-cad-RFP and VE-DEE-RFP loss at cell borders and down-regulation by both western blot and flow cytometry. The VE-KK-RFP border localization was slightly reduced, and levels of the protein were also decreased. In contrast, the VE-KKDEE-RFP mutant was fully resistant to MARCH2 mediated down-regulation. These data suggest that MARCH2 induced VE-cadherin downregulation depends primarily on juxtamembrane lysine residues with minimal dependency on the DEE constitutive endocytosis motif.

**Fig 4.**
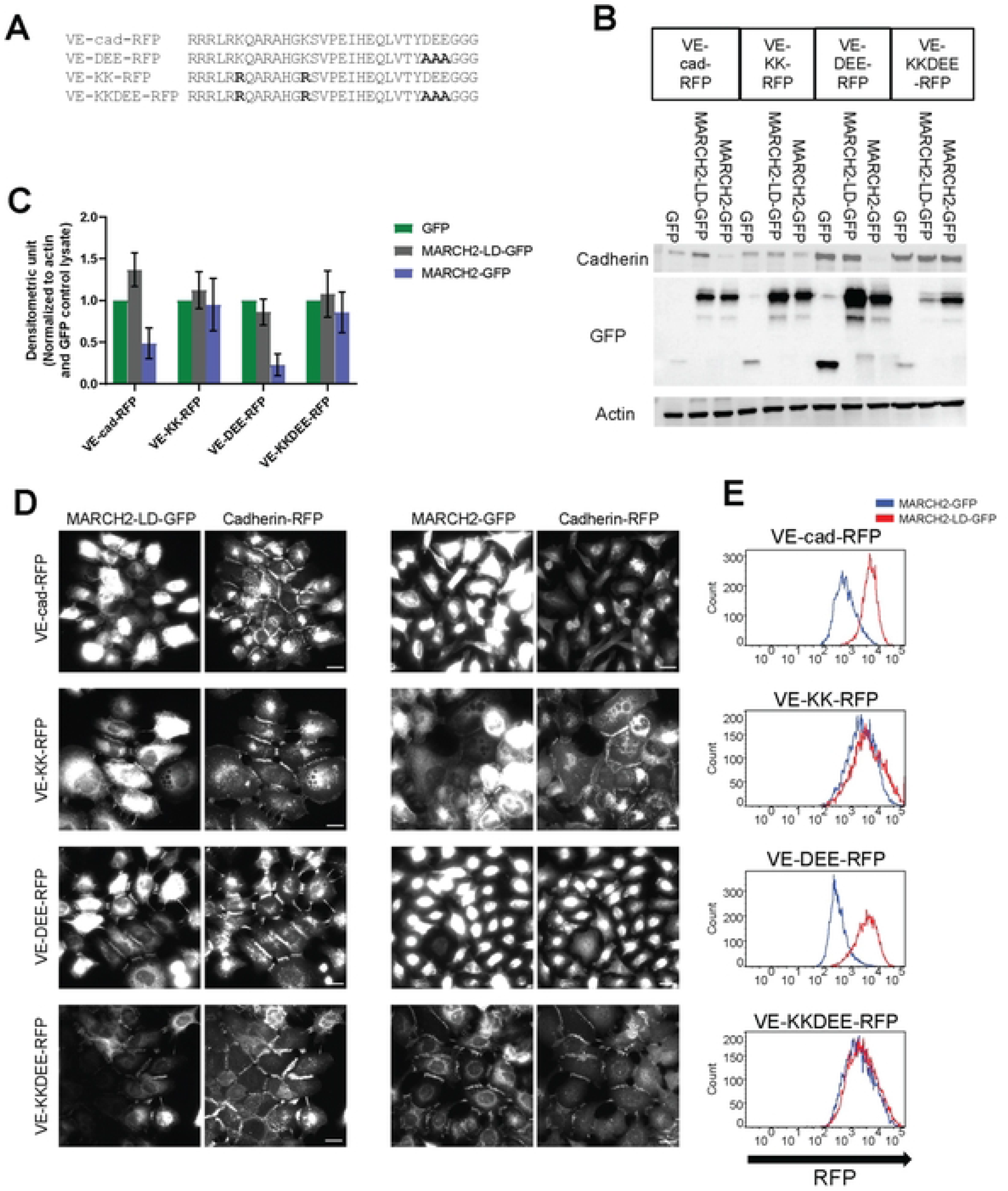
Juxta-membrane lysine residues are needed for MARCH-induced downregulation of VE-cadherin. (A) Juxta-membrane sequences of VE-cadherin and VE-cadherin mutants are shown with bolded **m**utation sites. (B) Western blot analysis of cells expressing various cadherin mutants. Proteins are detected using antibodies directed against VE-cadherin, actin or GFP antibody. (C) Band densities are presented as mean +/- SEM from 3 replicates after normalization with GFP transfected samples. (D) MARCH2-LD-GFP (left) or MARCH2-GFP (right) were expressed using adenoviral delivery systems in E-cadherin/P-cadherin null A431 cells expressing various VE-cadherin mutants. 24 hours after transfection, cells were fixed and processed for fluorescence microscopy. MARCH-GFP and MARCH2-LD-GFP are shown on left and cadherin mutants on right. Scale bar = 25 µm. (E). MARCH2-LD-GFP or MARCH2-GFP were transiently expressed in stable A431 cells expressing various RFP tagged VE-cadherin mutants and subjected to flow cytometry. Living GFP positive cells **we**re **counted after** filtering by FSC, SSC and GFP values. Vertical axis indicates counts of signals horizontal axis indicates cadherin-RFP signal intensity. Red line indicates MARCH2-LD-GFP transfected cells and blue line indicates MARCH2-GFP transfected cells.

Surprisingly, we observed that MARCH2-GFP colocalized at cell-cell borders with both the VE-KK-RFP and VE-KKDEE-RFP mutants, both of which are resistant to downregulation by MARCH2. Similarly, MARCH2-LD-GFP colocalized with WT VE-cadherin-RFP at cell borders (Fig 4C). These results suggest that MARCH2 and VE-cadherin interact directly or indirectly at the plasma membrane, and that failure to ubiquitinate the substrate traps the MARCH ligase-cadherin complex as a stable enzyme-substrate intermediate. We have previously shown that p120-catenin dissociates from classical cadherins during endocytosis [16], therefore, we tested the possibility that MARCH2 binding to VE-cadherin displaces p120-catenin. To assess this possibility, we expressed the various cadherin mutants in a cadherin null background using A431 cells deficient in both E-cadherin/P-cadherin (Fig S3 and [43]. p120-catenin failed to localize to junctions in the cadherin null cells as shown in Fig S3 but localized at cell borders in cells expressing the various cadherin mutants (Fig S4). As we previously reported, the DEE to AAA mutation reduces p120-catenin binding and mutants harboring the DEE mutation exhibited reduced p120-catenin at cell junctions (Fig S4). p120-catenin was diffusely distributed when VE-cad-RFP and VE-DEE-RFP were downregulated by MARCH2 (data not shown). On the other hand, p120-catenin remained at cell borders and colocalized with MARCH2 in cells expressing the VE-KKDEE-RFP mutant (Fig 5A). Quantitative assessment failed to reveal any effect of MARCH2-GFP on p120-catenin colocalization in cells expressing cadherins that were resistant to down-regulation, such as the VE-KK and VE-KKDEE mutants (Figs 5B, C). These results suggest that p120-catenin and MARCH2 binding to VE-cadherin can occur simultaneously, but that ubiquitination of the cadherin tail causes p120-catenin dissociation and VE-cadherin endocytosis.

**Fig 5.**
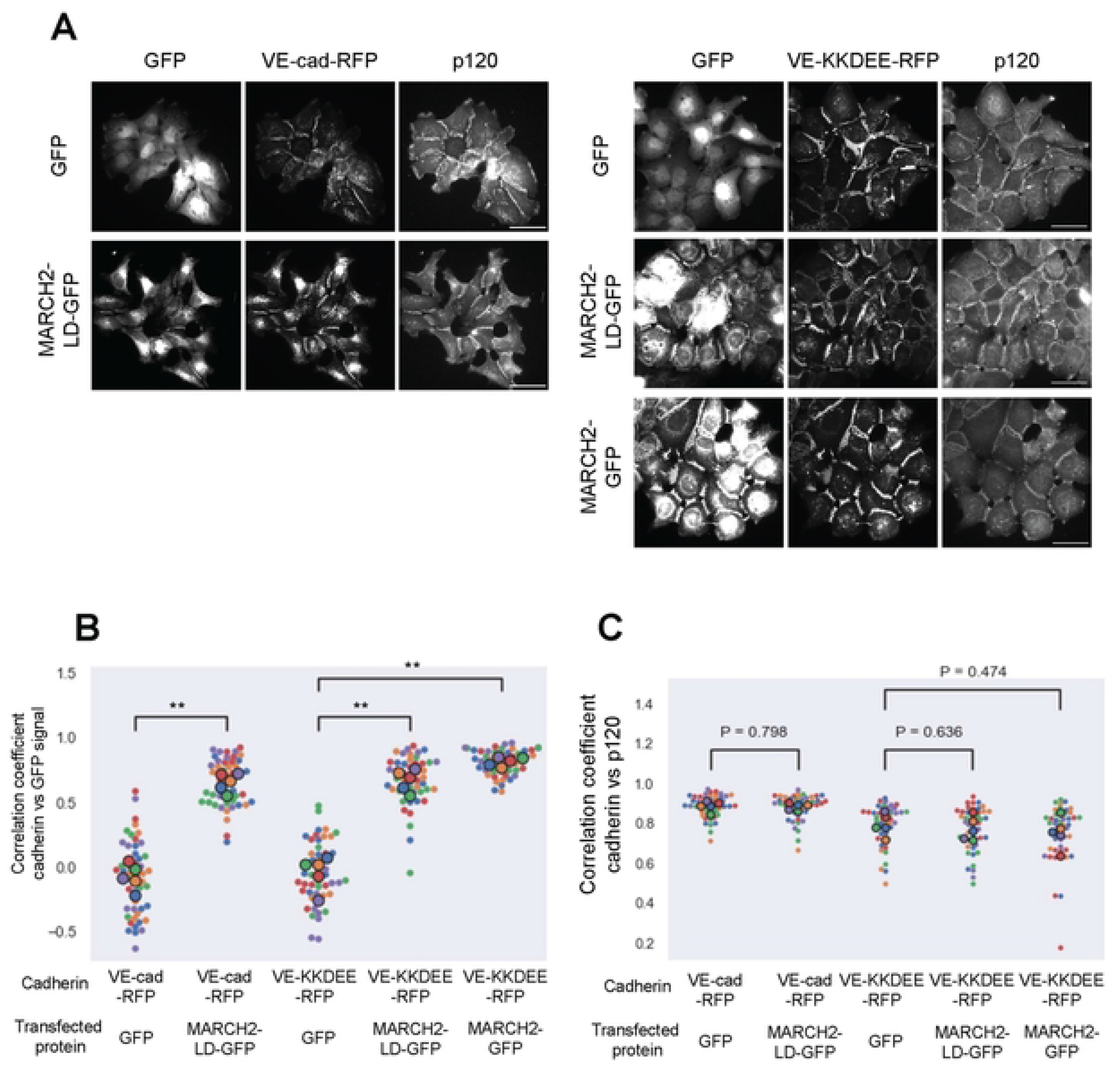
MARCH2 colocalizes at cell borders in cells expressing VE-cadherin mutants resistant to degradation. (A) GFP, MARCH2-LD-GFP, or MARCH2-GFP were expressed in E-cadherin/P-cadherin null A431 cells stably expressing VE-cad-RFP or VE-KKDEE-RFP. Scale bar = 50 µm. (B, C) Fluorescence intensities from 10 cell borders in 5 fields of view for each group were measured and used for calculating the Spearman’s Rank Correlation Coefficient (SRCC) between RFP and GFP (B) or GFP and p120-catenin (C). Smaller dots represent individual cell borders. Larger dots represent the average value from each field of view. Color represents data from the same field of view. **<0.01.

Collectively, the data presented here indicate that MARCH family ligases exhibit differential selectivity for cadherin family members. This selectivity is conferred predominantly by the cadherin transmembrane domain, which likely interacts with MARCH family ligases at the plasma membrane to drive cadherin endocytosis and degradation. The ligase then utilizes lysine residues in the cadherin juxtamembrane domain for ubiquitination to trigger internalization of cell surface cadherin. This MARCH induced cadherin ubiquitination pathway is distinct from cadherin ubiquitination by Hakai, a cytoplasmic ubiquitin ligase that induces E-cadherin downregulation in a phosphorylation dependent manner [24]. The phosphorylated residues needed for Hakai-mediated degradation of E-cadherin are not conserved in VE-cadherin, further indicating that cadherins are differentially regulated by a variety of E3 ubiquitin ligases.

It is particularly interesting that N-cadherin is resistant to every MARCH family ligase that we tested. N-cadherin often remains or is upregulated in epithelia and endothelia undergoing transitions to a more mesenchymal state [3, 15, 44-47]. These findings raise the intriguing possibility that MARCH family ligases might be involved in E(nd)MT-like processes during development or cancer progression that result in the loss of VE-cadherin or E-cadherin while preserving N-cadherin levels. This selectively of cadherin down-regulation would support a more migratory and mesenchymal phenotype. Further studies are needed to determine when MARCH family ligases are expressed and how they are regulating in various developmental and disease processes characterized by changes in cadherin expression profiles.

## Acknowledgements

We thank Dr. C. M. Grimsley-Myers and Dr. R. H. Isaacson for assistance with lentivirus production. We thank Drs. Mark Udey and Chuanjin Wu for providing cDNA reagents for various MARCH family ligases. We thank Nate Sheaffer and Joseph Bednarczyk from Penn State College of Medicine’s Flow Cytometry Core (RRID:SCR_021134) for assistance with flow cytometry analysis and cell sorting. This work was supported by grants from the National Institutes of Health (R01AR050501 and AR048266).

## Supporting information

**Fig S1. E-cadherin localization in MARCH-transfected cells.** A431 cells were transfected with MARCH family proteins and subjected to immunofluorescence with E-cad antibody then processed for fluorescence microscopy. Transfected cells are indicated with yellow arrowhead. Scale bar = 25 µm

**Fig S2. Cadherin chimeras localize to cell-cell junctions in A431 cells.** Localization of RFP tagged various cadherin chimeras were assessed in A431 cells expressing GFP. Scale bar = 50 µm

**Fig S3. Localization of cell adhesion molecules in A431 cells and A431 E-cadherin/P-cadherin null cells.** Wild type A431 cells or A431 cells lacking E- and P-cadherin [43] were processed for fluorescence microscopy to localize E-cadherin, N-cadherin, β-catenin or p120-catenin. The lack of β-catenin and p120-catenin at cell-cell contacts confirms lack of other classical cadherins. Scale bar = 50 µm

**Fig S4. Localization of VE-cadherin mutants lacking membrane proximal lysine residues.** E-cadherin/P-cadherin null A431 cells expressing various VE-cadherin mutants were analyzed by immunofluorescence for localization of the cadherin and p120-catenin in cells expressing GFP. Scale bar = 25 µm

## Material and methods

### Cells and plasmids

A431 E-cadherin and P-cadherin null cells were obtained as previously described. HUVECs were obtained from Lonza. VE-cadherin mutants were made as previously described [9]. N-cadherin cDNA was acquired from addgene (Plasmid #38153) [48]. N-cadherin and VE-cadherin cDNA sequences were subcloned into pLenti6/V5-DEST vector then cDNA sequences were modified to make mutants and chimeras. MARCH family proteins cDNA and ligase dead MARCH2 is obtained as described [9].

HEK 293A (QBI293A, Qbiogene, CA) were cultured in DMEM (Corning 10-13-CV) supplemented with 10% FBS (Sigma) and 1% Penicillin/Streptomycin (Sigma). HUVEC (Lonza or gift from Dr. Alejandro Adam, [49]) were cultured using Endothelial Cell Growth Medium 2 (ECGM2, PromoCell, DE). For all HUVEC experiments, cells were used between P4 and P8, and plated onto surfaces coated with 0.1% gelatin (Sigma) at a density of 100,000 cells/cm^2^. All cells were cultured at 37°C under 5% CO2.

### Cell culture and immunofluorescence

All cells were cultured at 37 degrees Celsius and 5% CO_2_. HUVECs were cultured in Endothelial Cell Growth Medium (Sigma-Aldrich) with Antibiotic-Antimycotic Solution (Sigma-Aldrich). All A431 cell lines were cultured in DMEM with 4.5 g/L glucose, L-glutamine and sodium pyruvate (Corning) and with 10% FBS (R&D systems).

For localization studies in Figs 1, 3, 4, 5, S1, S2, S3, and S4, cells were fixed with 4% PFA at room temperature. Nonspecific staining was blocked using 0.1% BSA. After fixation, cells were incubated with primary antibody for 1 hour at room temperature. After washing with PBS+ 3 times, cells were incubated with secondary antibody for 1 hour at room temperature. After washing with PBS+, coverslips were mounted using ProLong Glass Antifade Mountant (Thermo Fisher Scientific) and were incubated overnight. Immunofluorescence images in Figs 1, 3, 4, 5, S1, S2, S3 and S4 were captured using a Leica DMi8, a 63×/1.40 NA HC PL APO CS2 oil objective, a Leica QWF-S-T filter cube, and a Hamamatsu ORCA-Flash4.0 V3 (C13440-20C CL) Digital Camera using LAS X software version 3.6.0.20104. Antibody information is noted in Tables 1 and 2.

**Table 1.** Primary antibody information.

**Immunofluorescence**
| Antigen | Company | Catalog number | Concentration | Antibody species | Figure number |
| --- | --- | --- | --- | --- | --- |
| VE-cadherin | BD | 610252 | 1/250 | Mouse | Fig 1 |
| N-cadherin | BD | 610920 | 1/200 | Mouse | Figs 1 and S3 |
| p120 catenin | BD | 610134 | 1/100 | Mouse | Figs 1, 5, S3 and S4 |
| Phalloidin-Alexa Fluor594 | Thermo/Invitrogen | A12381 | 2 unit/ml |  | Fig 2A |
| VE-cadherin | Santa Cruz | sc-6458 | 1/500 | Goat | Fig 2A |
| p120 catenin | Santa Cruz | sc-1101 | 1/500 | Rabbit | Fig 2B |
| N-cadherin | BD | 610920 | 1/100 | Mouse | Fig 2B |
| E-cadherin | BD | 610182 | 1/500 | Mouse | Fig S1 |
| beta catenin | Sigma | c-2206 | 1/1000 | Rabbit | Figs S1 and S3 |
| E-cadherin | abcam | ab1416 | 1/100 | Mouse | Figs S3 |

**Western Blots**
| Antigen | Company | Catalog number | Concentration | Antibody species | Blocking | Figure number |
| --- | --- | --- | --- | --- | --- | --- |
| VE-cadherin | Santa Cruz | sc-6458 | 1/1000 | Goat | 5% Non-fat Dried Milk | Fig 2C |
| N-cadherin | BD | 610920 | 1/1000 | Mouse | 5%BSA | Fig 2C |
| VEGFR2 | Cell Signaling Technology | 2479 | 1/1000 | Rabbit | 5%BSA | Fig 2C |
| Tag(CGY)FP | Evrogen | AB121 | 1/1000 | Rabbit | 5% Non-fat Dried Milk | Fig 2C |
| Actin | Bio-Rad | 12004163 | 1/1000 | Human | 5% Non-fat Dried Milk | Fig 2C |
| GFP | Thermo/Invitrogen | A11122 | 1/2000 | Rabbit | 5% Non-fat Dried Milk | Fig 4 |
| beta-actin | Cell Signaling Technology | 8457 | 1/1000 | Rabbit | 5% Non-fat Dried Milk | Fig 4 |
| VE-cadherin | Beckman Coulter | A07481 | 1/500 | Mouse | 5% Non-fat Dried Milk | Fig 4 |

**Table 2.** Secondary antibody information.

| Name | Company | Catalog number | Concentration | Antibody species | Figure number |
| --- | --- | --- | --- | --- | --- |
| Alexa Fluor 555 anti-mouse IgG1 | Thermo/Invitrogen | A21127 | 1/3000 | Goat | Fig 1 |
| Alexa Fluor 647 anti-Goat IgG (H+L) | Thermo/Invitrogen | A21447 | 1/500 | Donkey | Fig 2A |
| Alexa Fluor 350 anti-Goat IgG (H+L) | Thermo/Invitrogen | A21081 | 1/500 | Donkey | Fig 2A |
| Alexa Fluor 594 anti-Mouse IgG (H+L) | Thermo/Invitrogen | A21203 | 1/500 | Donkey | Fig 2B |
| Alexa Fluor 350 anti-Rabbit IgG (H+L) | Thermo/Invitrogen | A10039 | 1/500 | Donkey | Fig 2B |
| HRP-conjugated anti-Goat IgG | Jackson ImmunoResearch | 805-035-180 | 1/10000 | Bovine | Fig 2C upper |
| StarBright Blue 700 anti-Rabbit IgG | Bio-Rad | 12004161 | 1/5000 | Goat | Fig 2C upper and lower |
| hFAB Rhodamine anti-Actin | Bio-Rad | 12004163 | 1/1000 | Human | Fig 2C upper |
| HRP-conjugated anti-Mouse IgG (H+L) | Jackson ImmunoResearch | 715-035-150 | 1/10000 | Donkey | Fig 2C lower |
| hFAB Rhodamine anti-Actin | Bio-Rad | 12004163 | 1/2000 | Human | Fig 2C lower |
| HRP-conjugated anti-Rabbit IgG (H+L) | Bio-Rad | 170-6515 | 1/3000 | Goat | Fig 4 |
| HRP-conjugated anti-mouse IgG (H+L) | Bio-Rad | 170-6516 | 1/3000 | Goat | Fig 4 |
| Alexa Fluor 647 anti-mouse IgG (H+L) | Thermo/Invitrogen | A21236 | 1/3000 | Goat | Figs 5 and S4 |
| Alexa Fluor 488 anti-mouse IgG2a | Thermo/Invitrogen | A21131 | 1/3000 | Goat | Fig S1 |
| Alexa Fluor 647 anti-mouse IgG (H+L) | Thermo/Invitrogen | A21236 | 1/1000 | Goat | Fig S3 |
| Alexa Fluor 555 anti-rabbit IgG (H+L) | Thermo/Invitrogen | A21428 | 1/1000 | Goat | Fig S3 |

For immunofluorescence studies in Fig 2, cells were grown in μ-slide 8 well plates (Ibidi). Cells were fixed for 30 minutes at 4°C with 4% paraformaldehyde in PBS (Thermo Fisher Scientific), briefly washed three times, then permeabilized for 15 minutes with 0.1% Triton-X 100 (Sigma). Nonspecific staining was then blocked for 1 hour using 5% donkey serum (Sigma). After blocking, cells were incubated with primary antibody (see Table) in 1% BSA (Rockland) for 2 hours at room temperature. After three 10 minutes washes, cells were incubated with secondary antibody conjugated with Alexa 350, 594, or 647 and/or Phalloidin-Alexa Fluor 594 (Thermo Fisher Scientific) in 1% BSA for 1 hour at room temperature. After three 10 minutes washes, fixed HUVEC cells labeled with GFP, Alexa Fluor 350, Alexa Fluor 594, and Alexa Fluor 647 in Ibidi µ-slide IbiTreat 8-well plates with TBS were imaged. Images were acquired with a Zeiss Axio Observer.Z1 microscope with Axiocam MRm camera. An Excelitas X-Cite 120LED Boost light source was used in combination with the Zeiss filter set 34 (BP 390/22, FT 420, BP 460/50), Zeiss filter set 38 (BP 470/40, FT 495, BP 525/50), Zeiss filter set 31 (BP 565/30, FT 585, BP 620/60, and Omega Optical QMAX-FRed (BP635/30, 660LP, BP710/80). Immunofluorescence images were captured using Plan-Apochromat 20x/0.8 M27 objective. Axiocam MRm camera was used with a 16-bit digitizer, 2x gain, and no binning. Acquisition was controlled with the Zeiss Zen 2.0 software. Antibody information is noted in Tables 1 and 2.

### Colocalization Analysis

For quantifying colocalization in Fig 5, images were analyzed with an ImageJ plugin, EzColocalization[50]. Briefly, each condition had five images, and 10 different cell borders (per image) were cropped for further analysis. Within EzColocalization, a minimum area threshold of 5 pixels was chosen to reduce spurious signals. ImageJ macros for cropping and running EzColocalization are available at https://zenodo.org/record/8147838.

### Flow cytometry

Cells were cultured in 6 well plate with 50% confluency and transfected by adenovirus. After 24 hours, cells were detached using TrypLE and resuspended in PBS-prior to flow cytometrywith a BD FACSymphony (BD Bioscience). Data analysis was performed using FlowJo (BD Biosciences). Dead cells were excluded by filtering forward scatter (FSC) and side scatter (SSC) value. GFP positive cells were filtered by comparing untransfected parental A431 cells.

### Western blot

Cells were grown in 24-well plates (BD) and lysed in 125 μl ice cold Laemmli buffer supplemented with Protease/Phosphatase inhibitor cocktail (Pierce). Lysates were heated 5 minutes at 95°C and stored at -20°C until processed. Thawed lysates were run on 7.5% Acrylamide gels with a 29:1 ratio Acrylamide:Bis using a Protean Tetra system (Bio-Rad) and Precision Plus markers (Bio-Rad). Gels were then transferred to 0.2 μm nitrocellulose (Bio-Rad) using the Criterion blotter (Bio-Rad). Proper transfer was verified by staining with Ponceau red, after which blots were washed, then blocked with 5% non-fat dry milk or BSA for 30 minutes. Blots were probed with primary antibodies overnight at 4°C in locker. The following day, blots were rinsed five times and washed for 30 minutes at room temperature, and then probed one hour at room temperature with bovine anti-goat HRP (Jackson) if a goat primary was used. Following this, blots were again rinsed and washed, and then probed with other HRP secondaries (Jackson), goat anti-mouse or goat anti-rabbit StarBrightBlue700 (Bio-Rad), and hFAB Actin Rhodamine primary antibody (Bio-Rad) for 1 hour at room temperature while protected from light. Blots were rinsed and washed, incubated with Clarity ECL reagent (Bio-Rad), and then imaged using a ChemiDoc MP system (Bio-Rad). Band density was measured using a Bio-Rad GelDoc imaging system (Bio-Rad Laboratories) and Image Lab analysis software (Bio-Rad Laboratories). Antibody information is noted in Tables 1 and 2.

### RT-PCR

HUVECs were cultured in 6 cm plates as described above. Cells were then scraped from plates and collected by centrifugation. RNA was purified from HUVECs by using RNeasy Mini kit (QIAGEN). cDNA was synthesized using the SuperScript III First-Strand Synthesis System (Invitrogen) and PCR was performed by using Platinum Taq DNA polymerase kit (Thermo Fisher scientific). The following primers were used: ACTB-Forward 5’-AGGATTCCTATGTGGGCGAC-3’; ACTB-Reverse 5’-ATAGCACAGCCTGGATAGCAA-3’; MARCH1 -Forward 5’-TCCGGAAGTGGGAGAAACT-3’; MARCH1-reverse 5’-CTCCTGGGCATTTGGTCTGT-3’; MARCH2-Forward 5’-GCCTCGACCCCTCACAGA-3’; MARCH2-reverse 5’-AATGAGATCAGGGTGCCTGC-3’; MARCH3-Forward 5’-ACTTGGCACTTAGAACCGCA-3’; MARCH-reverse 5’-CTCAGCCACTCCACTAACGG-3’; MARCH4-Forward 5’-CGTGGTGTGCATAGGTCTCAT-3’; MARCH4-reverse 5’-GGCCAGGTTCAGCCTGTATT-3’; MARCH5-Forward 5’-CCGCCTCTCAGTGCTATTGT-3’; MARCH5-reverse 5’-ACAAACGCAATTCCACCCAAG-3’; MARCH6-Forward 5’-GCCGGATACTTGCTGGATCT-3’; MARCH6-reverse 5’-CACAGGGAAAGCAACAGCAC-3’; MARCH8-Forward 5’-TCTCCTCAGCAGTACGGTCA-3’; MARCH8-reverse 5’-AGCAACATTTCTAAGGCTGGGA-3’; MARCH9-Forward 5’-GGCTCCGCATGTTTCTGAAC-3’; MARCH9-reverse 5’-GATGAGGCCTATGCAGACGA-3’; MARCH11-Forward 5’-CGGAGCAGGGTGAGTTGTTGA-3’; MARCH11-reverse 5’-CGGCTGCTTTCTTCGATGTC-3’.

### Adenovirus production

High titer adenoviruses to EGFP, MARCH2-GFP, MARCH2-LD-GFP, and MARCH4-GFP were produced in HEK 293A, concentrated by cesium chloride gradient, and then buffer exchanged by dialysis. Briefly, expression constructs were transferred to an entry vector and then shuttled to pAdCMV/V5/DEST destination vector via Gateway cloning LR reactions, sequence verified, then transfected and propagated using 293A, as described in the ViraPower Adenoviral Expression System manual (Thermo). Amplified adenoviruses were concentrated by cesium chloride gradient using a Beckman-Coulter Optima XL-100K ultracentrifuge equipped with SW41Ti rotor, by two successive 25000rpm runs over layered gradients using equal portions 1.33 and 1.45g/cm^3^ cesium chloride. Viral bands were collected and then dialyzed using a Tube-O-Dialyzer according to manufacturer’s instructions (G Biosciences) to buffer exchange to 10 mM Tris, pH 8, 2mM MgCl_2_, 5% Sucrose solution. Purified adenoviruses were then aliquoted and stored at -80°C.

### Isolating cell lines with cadherin

Lentivirus are produced by pLenti6/V5 DEST lentiviral delivery system (Thermo Fisher Scientific). A431 cells or E-cad P-cad null A431 cells were transduced using lentivirus in the medium containing 10 μg polybrene. 24 hours after transduction, cells were cultured in 10 cm plates and incubated to 70% confluence. Cells were detached by TrypLE Express Enzyme (Thermo Fisher scientific) and subjected to flow sorting by Aria SORP high-performance cell sorter (BD Biosciences). RFP positive cells were sorted and subcultured for recovery. RFP positive cells were then plated at low density, and RFP positive clones were isolated using cloning cylinders. Clones of interest were then expanded and characterized for fluorescence intensity, fluorescence localization, and cell morphology. Clones with normal and homogenous cadherin expression then were used for experiments.

### Endothelial Monolayer Permeability

Changes in endothelial monolayer permeability were assessed using Electric Cell-substrate Impedance Sensing (ECIS; Applied BioPhysics, [51], as indicated by changes in electrical resistance, as described previously [52]. Resistance correlates with barrier function, such that increasing resistance represents a tightening of the endothelial barrier, while a decrease represents a loss of barrier, and increased permeability. For ECIS experiments, HUVEC were seeded onto ECIS 8W10E PET culture ware (Applied Biophysics), and readings were taken every 5 minutes at single frequency 4000Hz. Cells were seeded at confluence and junctions were allowed 72 hours to mature, at which point cells were transduced with adenovirus to express MARCH proteins and followed for an additional 24 hours of monitoring.

